# The nuclear lamin network passively responds to both active or passive cell movement through confinements

**DOI:** 10.1101/2024.09.27.615360

**Authors:** Sirine Amiri, Inge Bos, Etienne Reyssat, Cécile Sykes

**Author notes:** These authors contributed equally to this work (Sirine Amiri, Inge Bos).

## Abstract

Physical models of cell motility rely mostly on cytoskeletal dynamical assembly. However, when cells move through the complex 3D environment of living tissues, they have to squeeze their nucleus that is stiffer than the rest of the cell. The lamin network, organised as a shell right underneath the nuclear membrane, contributes to the nuclear integrity and stiffness. Yet, its response during squeezed cell motility has never been fully characterised. As a result, up to now, the interpretations on the lamin response mechanism are mainly speculative. Here, we quantitatively map the lamin A/C distribution in both a microfluidic migration device and a microfluidic aspiration device. In the first case, the cell is actively involved in translocating the nucleus through the constriction, while in the second case, the cell behaves as a passive object that is pushed through the constriction by an external pressure. Using a quantitative description of the lamin shell response based on mass conservation arguments applied on the fluorescence signal of lamin, we show that in both cases of migration and aspiration, the response of the lamin network is passive. In this way, our results not only further elucidate the lamin response mechanism, but also allow to distinguish this passive deformation response from other active responses that may occur when the nucleus is squeezed through constrictions.

## 1 Introduction

The migration of cells through constrictions is essential for a wide variety of biological processes like embryonic development,[1] stem cell maturation,[2] immune response[3] and cancer metastasis.[4] The nucleus forms the limiting factor for this confined cell migration because it is the largest and stiffest organelle of the cell.[5, 6]

The translocation of the nucleus through a constriction is a complex operation: multiple proteins are involved, ranging from the proteins that regulate the compaction of chromatin in the nuclear interior to the proteins that are responsible for the cytoskeleton forces in the cytoplasm,[7] and different processes occur at the same time. Some of these processes are a purely passive response to the deformation of the nucleus. Other processes involve an active response of the cell. For example, the cell can use mechanotransduction to convert the mechanical stimuli into biochemical signals.[8] The fact that multiple processes occur simultaneously makes it challenging to distinguish the individual effects. Separating these effects becomes even more difficult due to the variety of proteins that is involved and the possible interactions between these proteins.

One of the protein families that play an essential role in the nucleus translocation are the nuclear lamina. The nuclear lamina family is divided in A-type lamins (lamin A and C) and B-type lamins (lamin B1 and lamin B2). Together, these lamin types form a stable intertwined meshwork situated directly beneath the nuclear envelope.[9, 10] In this way, the lamina contribute to the nuclear integrity (both A-type and B-type lamins) and the nuclear stiffness (regulated by A-type lamins only).[11, 12] Perturbations in lamin gene expression or translation can give rise to a range of diseases, collectively termed laminopathies.[13] Notably, the decreased stiffness of the lamina, resulting in reduced nuclear resistance to cytoskeletal forces exerted from outside the nucleus, has been implicated as a potential factor in certain types of heart failures associated with laminopathies.[14] Furthermore, dysregulation of lamin proteins, either upregulation or downregulation, is observed in various cancer types, although its implications for cancer cell motility remain inconclusive.[15–17]

When cells migrate through constrictions the intensity of fluorescently labelled lamin increases at the sites where the nucleus is compressed.[4, 18, 19] It has been postulated that this intensity increase is a passive response and might be due to buckling of the lamin network.[18–21] However, so far, this lamin intensity increase has never been fully quantified. As a result, this interpretation remains rather speculative.

In this paper, we quantify the lamin A/C distribution in both an aspiration device[22] and a migration device.[18] In particular, we introduce a new method to map the nuclear protein distribution. In addition, we provide a quantitative description of the lamin response based on mass conservation. Together, our results not only prove that the lamin network deforms passively, but also aid to separate this passive deformation response from any other effects that occur during confined cell migration. In this way, our findings can assist to further elucidate the complex process of confined cell migration.

## 2 Experimental results

### 2.1 Nuclear trajectories and deformations

Nucleus deformation is observed in two different situations using 5 µm-high microfluidic devices: i) cells spontaneously *migrate* through constrictions, ii) cells are *aspirated* into small channels.

The *cell migration* device [18] consists of a series of pillar structures realising constrictions of different widths (*w* =15, 5, 3 and 2 µm) (Fig. 1.a); cells spontaneously migrate from one side of the device to the other (see subsection 5.3). The *cell aspiration* device [22] is made of 3 µm wide rectangular channels (Fig. 1.a’); cells are aspirated through the channels by applying a controlled pressure difference (see subsection 5.4).

**Figure 1.**
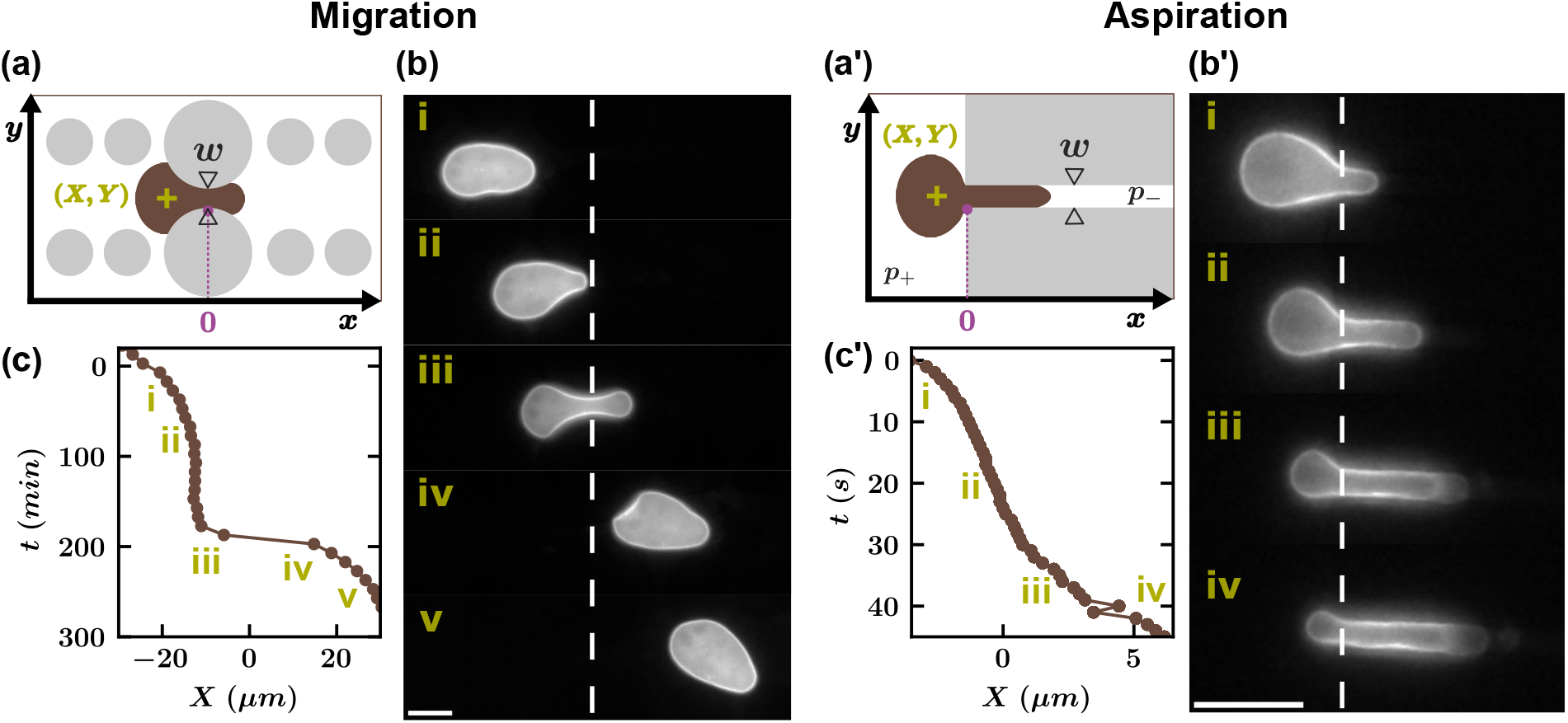
Nucleus trajectories during migration and aspiration. – (a) Top view of the pattern used in the *migration* device. It is composed of PDMS pillars of several widths in order to make three types of constrictions : 5, 3 and 2 µm wide and a control channel of 15 µm. The height of the pillars is *h* = 5*µm*. (a’) Top view of the pattern used in the *aspiration* device. A pressure difference Δ*p* ∼ 6900 Pa drives cells through the 3 µm wide and 5 µm high channel. (*X, Y* ) (*green cross*) is the barycenter of the nucleus (*brown*), used as a proxy for its position. (b-c) Example of the position of a MEF nucleus against time during *migration* through a 3µm wide constriction and corresponding epifluorescence mCherry images for selected time points (i to v). (b’-c’) Example of the position of a MEF nucleus against time during aspiration through a 3µm wide channel and corresponding epifluorescence mCherry images for selected time points (i to iv). White dashed lines indicate where the origin of the *x*-axis is. Scale bars: 10µm.

The nuclear lamin shell is imaged by the endogenous lamin fluorescent signal in CRISPR-engineered Mouse Embryonic Fibroblasts (MEFs) expressing lamin A/C with an mCherry tag (see subsection 5.1). The fluorescence of lamin A/C allows for a clear detection of the nucleus shape during migration and aspiration, as shown in Fig. 1b,b’. The nucleus position (*X, Y* ) is defined as the surface geometrical barycenter. For the migration device, the origin of the *x*-axis is set in the middle of a given constriction defined by the two pillars (see subsection 5.7). For the aspiration device, the origin of the *x*-axis is set at the entry point of the constriction channel. In both devices, as a nucleus travels the constriction, it elongates in the direction of motion and undergoes transverse confinement.

The characteristic travelling time of nuclei across constrictions is several hours for spontaneous *migration* whereas it is tens of seconds for forced *aspiration*. The *migration* trajectories *X*(*t*) have a sigmoidal shape: the nucleus slows down when entering the constriction, and it accelerates when it is expelled from the constriction (Fig. 1c). In contrast, the *aspiration* trajectories *X*(*t*) appear smoother: the nucleus is transported continuously through the channel, as illustrated in Fig. 1c’.

Even though the trajectories and time scales are different between *migration* and *aspiration*, we observe a similar lamin A/C fluorescence increase at sites where the nucleus is most deformed, as displayed Fig. 1b and Fig. 1b’. During *migration*, the fluorescence is at highest on the sides of the nucleus when it is in the middle of the constriction (iii), whereas the fluorescent signal is homogeneous right before (i) and right after (v) the constriction (Fig. 1b). During *aspiration*, the fluorescence is stronger at the front sides of the nucleus, where it contacts the channel borders (ii-iv), compared to the initial position of the nucleus at the channel entry (i) (Fig. 1b’).

### 2.2 Fluorescence intensity profiles along the nucleus contour

To further address the similarity in lamin A/C fluorescence increase in both the *aspiration* and *migration* process, we set out to quantify lamin A/C fluorescence along the nucleus contour. We define the nucleus contour coordinate *s* and measure the lamin A/C fluorescence intensity as a function of *s*. In order to compare the profiles of different time points and nuclei with each other, we normalise the fluorescence intensity *I* with respect to the average intensity in the contour, and we normalise the nucleus contour coordinate *s* with respect to the nucleus perimeter *L*. The back of the nucleus is at *s* = 0 and the contour is oriented counter clockwise (Fig. 2).

**Figure 2.**
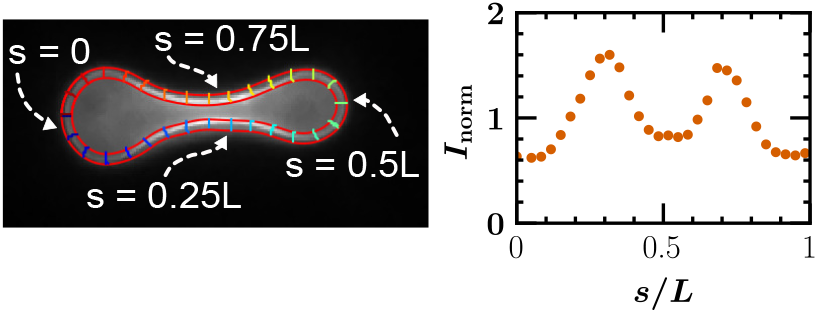
Example of the determination of the lamin A/C fluorescence intensity profile as a function of the nucleus contour coordinate *s. Left* Typical mCherry image of a MEF nucleus translocating through a 2 µm wide constriction. The nucleus contour is taken as a 1 µm wide band at the edge of the nucleus. The contour is divided into 30 segments along the curvilinear abcissa *s* and the fluorescence intensity of each segment is measured. *Right* Normalised lamin A/C fluorescence intensity as function of *s/L* for the image shown on the left.

For each position X of the nucleus we average the fluorescence intensity profiles of several experiments.

In this way, we can follow how the averaged intensity distribution evolves when the nucleus deforms.

For the *migration* device, we focus on five reference positions of the nucleus, the central one corresponding to the nucleus positioned at the center of the constriction (*X* = 0 *µm*), as sketched in Fig. 3a, *top*. Peaks are observed in intensity profiles for the three central nucleus positions (*X* = −10, 0, 10 *µm*) (Fig. 3a, *bottom*) consistent with qualitative observations of Fig1b. The lamin intensity increase is distinctly more pronounced at *X* = 0 *µm*. At *X* = −20 *µm* and *X* = 20 *µm*, when the nucleus is not in contact with the pillars, the lamin intensity signal is constant along the entire perimeter.

**Figure 3.**
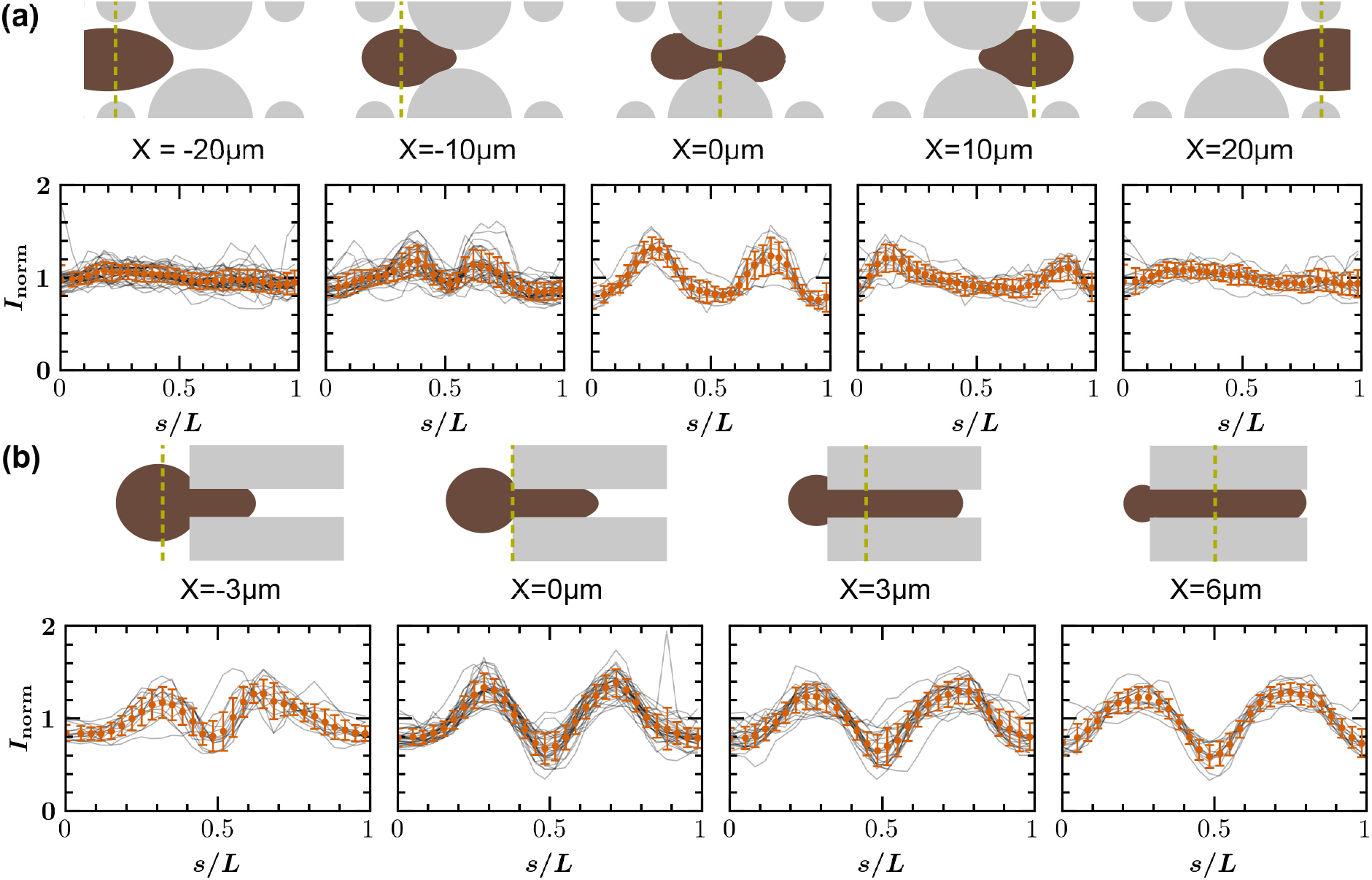
Lamin A/C fluorescence intensity profiles at different key nucleus positions during migration and aspiration – Sketches representing a nucleus (*brown*) crossing a 3 µm constriction in the migration device (a)-*top*, or a 3 µm wide channel in the aspiration device (b)-*top* at different key nucleus positions. Green dashed lines represent where the barycenter of the nucleus (*X*) is localised. (a) and (b)-*bottom* Lamin A/C intensity signal measured along the curvilinear abscissa *s/L* at the corresponding nucleus positions. For each position *X*, we average the signal corresponding to a number *N* of cells for which we possess images at *X ±* 5*µm* in the migration experiment and at *X* ± 0.3*µm* for the aspiration experiment. Grey lines are individual intensity profiles. Mean values are represented with circular symbols. Error bars represent standard deviation. Intensities are normalised (see subsection 5.7). For (a)-*bottom*, from left to right : N=38, N=34, N=12, N=15 and N=14. For (b)-*bottom*, from left to right : N=14, N=30, N=23 and N=14.

For the *aspiration* device, we define four key positions, the second one corresponding to the nucleus barycenter coinciding with the entrance of the channel as schematised Fig. 3b, *top*. Intensity peaks are the lowest at *X* = −3*µm*, when the nucleus is only slightly engaged in the channel (Fig. 3b, *bottom*).

The intensity profiles allow to quantitatively compare the lamin A/C intensity increase of the *migration* and *aspiration* process. In both cases, the peaks have a similar magnitude when the nucleus is in the middle of the constriction, suggesting that the similarity in fluorescence increase between the two processes is also quantitative. We note that the increase in lamin A/C intensity of the other nucleus positions is more difficult to compare directly due to the different geometry of the two devices.

### 2.3 Shift of the fluorescence peaks during nucleus translocation

In both the *migration* and *aspiration* devices, we observe two peaks that increase and move away from each other before *X* = 0*µm* (Fig. 3). In the *migration* device, for *X >* 0*µm*, the two peaks continue to move away from each other (i.e. localise at the back of the nucleus) and decrease in amplitude until they disappear when the nucleus exits the constriction. In the *aspiration* device, the peaks get wider as the nucleus further engages in the channel.

The location of the fluorescence peaks along the curvilinear abscissa *s/L* is determined for each position *X* of every nucleus (see details in subsubsection 5.7.2). In this way, we can plot the average peak position as function of the nucleus position *X* (Fig. 4).

**Figure 4.**
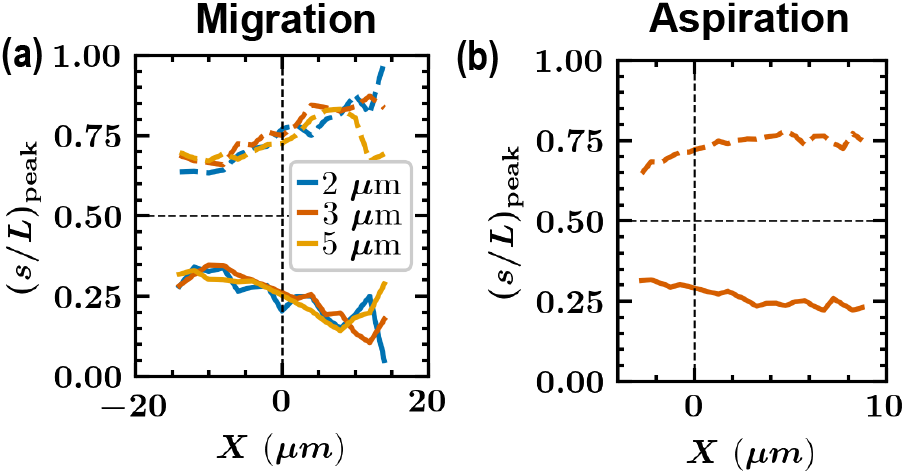
Shift of the average lamin A/C fluorescence peak position (*s/L*)_peak_ during migration and aspiration. – Average position of the fluorescence intensity peak as a function of nucleus position for different constriction sizes in the migration device (a) and in the aspiration device (b). Coloured solid lines (resp. dashed lines) correspond to the average position of the first peak (resp. second peak). Black dashed lines act as guides for the front of the nucleus (*s/L* = 0.5) and the position *X* = 0 *µ*m.

In the *migration* device, the peaks shift from the front to the back of the nucleus as it moves through the constriction. Hence, the first and second peak curves are symmetrical relative to the axis *s/L* = 0.5 (Fig. 4). Note that peak displacement does not significantly depend on the size of the constriction (2,3 and 5µm). The first peak position is symmetric around the point (*X* = 0, *s/L* = 0.25). Likewise, the second peak position is symmetric around the point (*X* = 0, *s/L* = 0.75)). From this symmetry around the central point *X* = 0, we conclude that peak position does not depend on the direction of motion, but only on the distance to the constriction center.

In the *aspiration* device, the peaks also shift backwards, but this shift is less pronounced than in the migration device, and further in the migration device the peak position is barely changing. Instead, it seems to have reached a constant position around (*s/L*)_peak_ ≈ 0.25 for the first peak and (*s/L*)_peak_ ≈ 0.75 for the second peak. The difference between the aspiration and migration device can be explained by their different geometry: the constriction is longer in the aspiration device and therefore the whole nucleus fits inside the constriction. Since the aspiration channel has a constant width, the most confined part of a well-engaged nucleus is its initially widest region, corresponding to (*s/L*)_peak_ = 0.25 and (*s/L*)_peak_ = 0.75. The nucleus often ruptures before exiting the aspiration channel. Therefore, the point where the nucleus exits the channel is not taken into account in the analysis. As a result, we do not have data points where the compression is focused at the back of the nucleus and hence we do not observe peak positions located at the back of the nucleus for the aspiration device.

Altogether these observations reveal that the hyperfluorescence of lamin A/C is correlated with nucleus deformation and not to the active motion of the nucleus.

### 2.4 Lamin A/C mobility

A possible explanation for the increase in lamin fluorescence is that lamin is recruited to the sites where the nucleus is constrained. Earlier measurements have shown that the lamin A/C in the lamina network is immobile on the time scales of both the aspiration and migration experiments.[19, 23, 24] However, the nucleoplasmic lamin A/C is much more mobile[23–25] and could thus be rapidly recruited to the compression sites. To test whether this is the case, we performed fluorescence recovery after photobleaching (FRAP) experiments where we bleached a part of the nucleus that is inside a constriction of the migration device (Fig. 5). Although there is indeed some initial fast recovery due to the nucleoplasmic lamins, this recovery is insufficient to regain the hyperfluorescence at the compression site. In fact, we do not observe any difference in lamin fluorescence intensity between the compression site and other positions in the nucleoplasm. Therefore, we conclude that the increase in lamin A/C fluorescence at the compression sites is not the result of the recruitment of lamin A/C from elsewhere in the nucleus.

**Figure 5.**
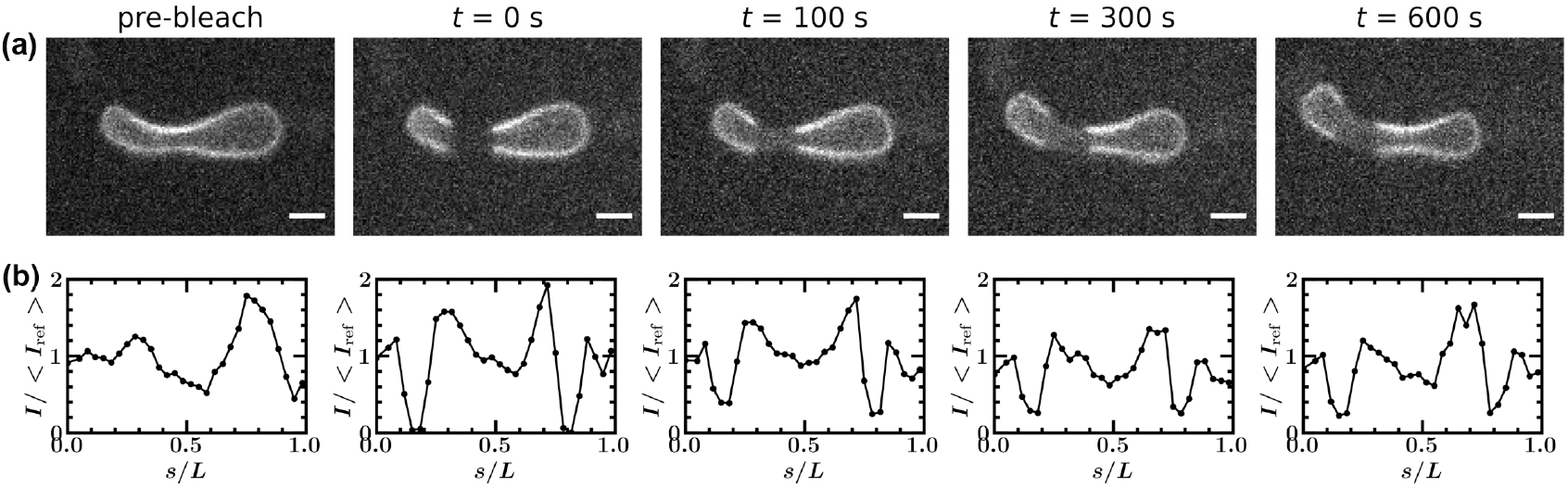
FRAP experiment of the lamin shell of a nucleus moving through a 3 *µ*m constriction. (a) Example images of the nucleus before bleaching (pre-bleach) and different time points after bleaching (bleaching is at t = 0 s). Scale bar: 5 *µ*m. (b) Corresponding fluorescence intensity profiles of the nucleus contour. The intensity profile is normalised by the average intensity in the contour before bleaching *< I*_ref_ *>*.

## 3 A minimal model

Altogether, our experimental results suggest that the nuclear lamina behaves as a passive surface. Since we use a CRISPR-modified cell line to mark lamin A/C, we know that we exclusively observe the fluorescence of endogenous lamins. Assuming that the fluorescent signal is proportional to the quantity of lamins, we now provide a minimal geometrical model to account for the variations in lamin fluorescence intensity along the contour of the nucleus. Mass conservation implies that the increase (resp. decrease) of fluorescence per unit area corresponds to an effective compression (resp. extension) of the lamin shell. We propose a minimal model that is independent of the detailed mechanism of lamin shell stretching or compression, as, for example, compression could be due to actual compression or to the wrinkling of the shell.

Let us first consider the geometry of the migration device. Before entering the constriction, the nucleus is not yet deformed and its shape is approximated by an elliptical pancake of length 2*a*_0_ = 20 *µ*m, width 2*b*_0_ = 12 *µ*m and height *h* = 5 *µ*m, with an associated total surface area *S*_0_ = 2*πa*_0_*b*_0_ + *ph*, where *p* is the perimeter of the undeformed nucleus contour. In agreement with experimental observations (Fig. 1(b)i and Fig. 3(a) left), the lamin fluorescence around this undeformed nucleus *I*_*ndef*_ is assumed to be homogeneous. Once in the constriction, nuclei are slightly elongated in the direction of motion while they are squeezed laterally. We compute the resulting change in nucleus contour and lamin signal in two steps. The first step consists in taking into account the longitudinal stretching of the nucleus that is observed experimentally, by defining a virtual stretched state of the nucleus. The maximal relative elongation *α*_*max*_ is reached at the center of the constriction, i.e. for *X* = 0 *µ*m (we measure experimentally the average value of *α*_*max*_ = 1.22, 1.12, 1.11 for *w* = 2, 3, 5 *µ*m respectively). For any position *X* of the nucleus, we use a parabolic interpolation *α*(*X*) = 1 + *α*_*max*_ (1 − (*X/X*^*\**^)^2^) where *X*^*\**^ is the position at which the nucleus first contacts the constriction (*X*^*\**^ = 15.9, 15.12, 13.29 *µ*m for *w* = 2, 3, 5 *µ*m respectively). To this virtual stretched state corresponds a change in total surface area of the nucleus which is *S*_*i*_ = 2*παa*_0_*b*_0_ + *p*_*d*_*h*, where *p*_*d*_ is the perimeter of the virtual elongated nucleus contour. This intermediate step induces a homogeneous decrease of the lamin signal by a factor *S*_0_*/S*_*i*_. In a second step, we squeeze laterally any section of the deformed nucleus which is wider than the constriction, giving rise to the final constricted shape of the nucleus. The resulting local increase of the fluorescence signal is computed by preserving the integrated fluorescence signal of a constant-*x* slice of the nucleus. A slice of initial width *D* and total length 2*D* + 2*h* is squeezed to a compressed width *d* and total length 2*d* + 2*h*. The final fluorescence signal reflects the two-step process so that *I*_*def*_ obeys:

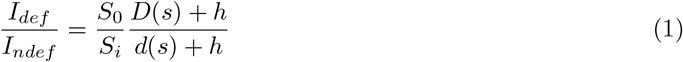

where *D*(*s*) and *d*(*s*) vary along the boundary of the nucleus, i.e. with *s* (Fig. 6). Comparing predictions made by the above equation with experimental intensity profile measurements reveal that this minimal model quantitatively accounts for the amplitude and position of the fluorescence peaks (Fig. 7, Fig. S1).

**Figure 6.**
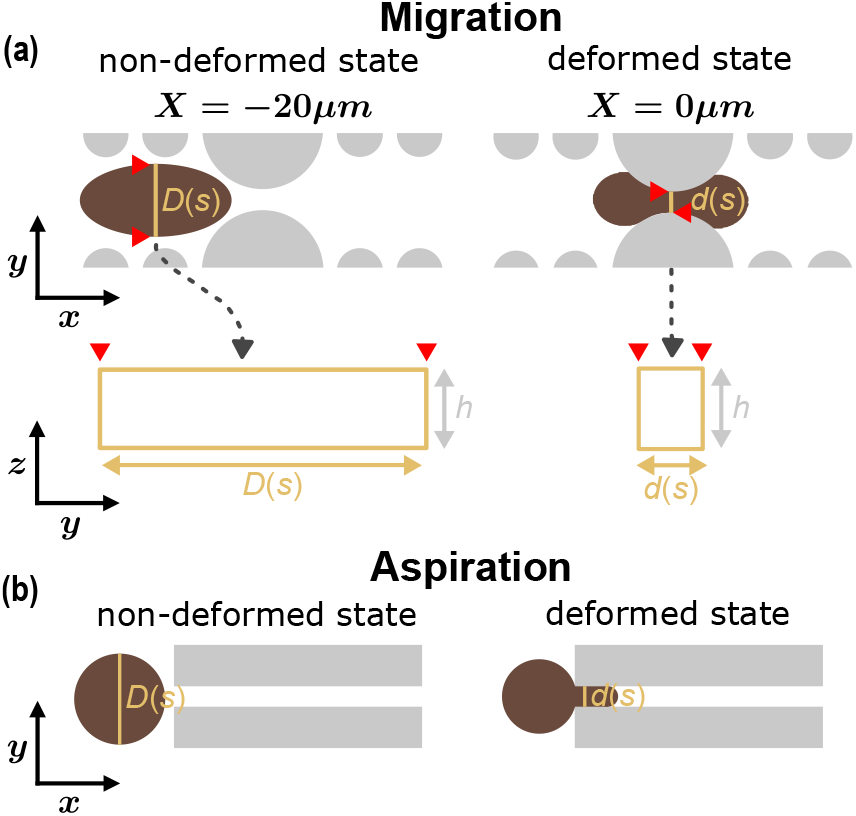
Sketches of the cross sections considered in both the migration and aspiration experiments. **(a)** *Top:* Representation of the *non-deformed* state at *X* = −20*µm*(*left* ) and a *deformed* state at *X* = 0*µm* (*right* ) of the nucleus (*brown*) in the (*x, y*) plane of the constrictions (*grey* ) in the migration experiment. *Bottom:* Representation of the orthogonal section (*yellow* ) approximated as a rectangle in the *non-deformed* (*right* ) and *deformed* (*left* ) state of the nucleus in the (*z, y* plane). Red arrows represent the location of the accessible lamin intensity in the migration experiment. **(b)** Representation of the nucleus in the aspiration experiment in a *non-deformed* state (*left* ) and a *deformed* state (*right* ) in the (*x, y*) plane.

**Figure 7.**
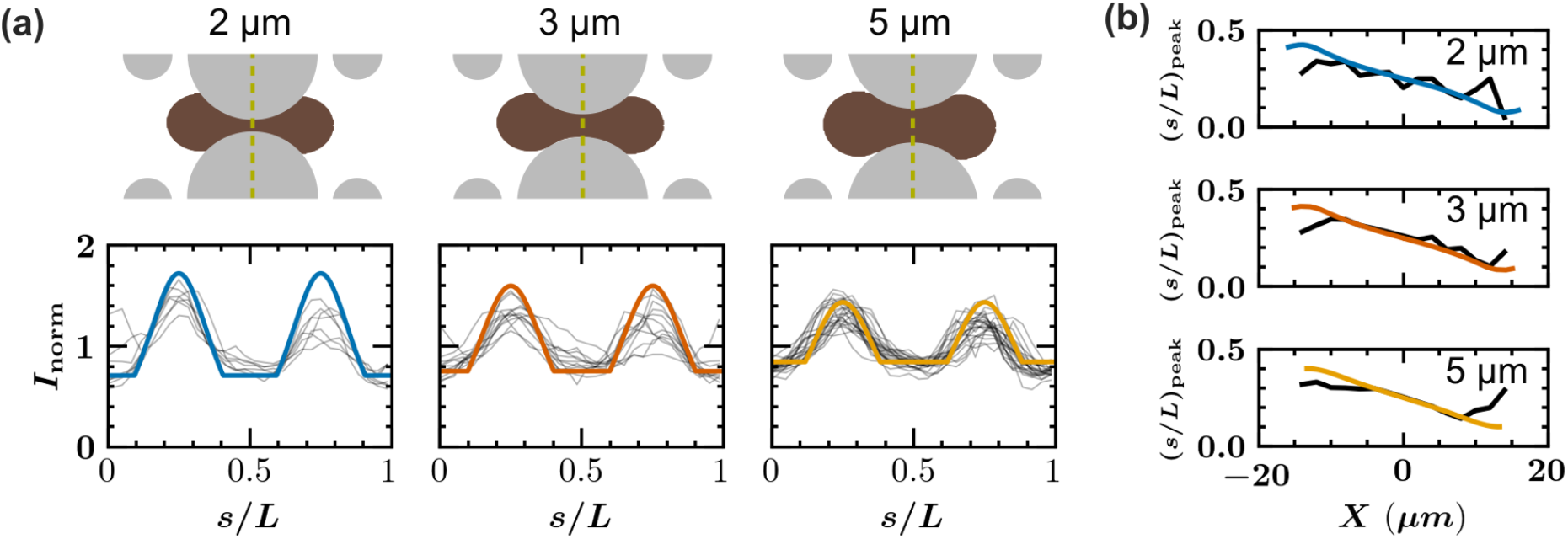
Comparison of the experimental lamin A/C spatial distribution to the minimal model predictions in the migration experiment. (a)-*top*: Sketches representing a nucleus (*brown*) at *X* = 0*µm* position for different constriction sizes. (a)*bottom*: Lamin A/C intensity signal measured along the curvilinear abscissa *s* at *X* = 0*µm*. For each position *X*, we plot individual intensity profiles (*grey lines*) of a number *N* of cells for which we possess images at *X* = 0 *±* 5*µm*. On top is plotted the minimal model prediction (*bold coloured line*). From left to right : N=7, N=12 and N=23. (b) Position of the fluorescence intensity peak (*s/L*)_peak_ as a function of nucleus position for different constriction sizes in the migration device. Coloured lines correspond to the model prediction and black lines correspond to the experimentally measured average position of the first peak.

A similar attempt is made at describing the fluorescence data from nuclei in the aspiration device. The geometry of the model is simplified, and the device is assumed to be cylindrical tube connected to a semi-infinite chamber. For the cylindrical tube diameter, we took the diameter for which the circumference of the cylinder *πd*_*c*_ roughly equals the circumference of the aspiration tube 2*d* + 2*h*, which gives *d*_*c*_ = 5 *µ*m. The nucleus, initially in the non-confined part, is assumed to be a sphere of radius *R*_0_ = 4 *µ*m. Once the nucleus is inserted in the channel, its shape is described in three sub-units: (i) a hemisphere at the front, with a diameter equal to that of the channel, (ii) a cylindrical part along the wall of the channel and (iii) a spherical cap at the back which is outside the channel. The arclength *L* from the back to the front of the contour is assumed to be conserved, *L* = *πR*_0_. This is a simplification because in experiments the nucleus stretches inside the channel resulting in an increase of the arclength with a factor ∼ 1.4. As earlier, the lamin signal intensity in the unconfined state is assumed to be homogeneous. We consider the part of the lamin shell initially at a distance *s* measured along the surface from the rearmost point of the nucleus. This material is assumed to remain at distance *s* from the rear at all times. Once the nucleus enters the channel, it thus undergoes an effective compression, from which we compute the predicted variation in lamin fluorescence. As seen from Fig. 8a and Fig. S2, this prediction agrees well with the experimentally measured intensity profiles. The differences in profile shape can be explained by the fact that we simplified the nucleus geometry in the model resulting in sharper transitions in the intensity profile predicted by the model. In addition, we ignored the stretching of the nucleus inside the device. Stronger stretching of the front of the nucleus might explain why the experimental front intensity is lower than the modelled front intensity. The simplification of nucleus geometry and neglecting the variability in nucleus shape might also account for the sharper transition in the peak position in the model prediction than in experiments (Fig. 8b). Still, the model is able to capture the experimental plateau of the peak position around (*s/L*)_peak_ ≈ 0.25 (Fig. 8b).

**Figure 8.**
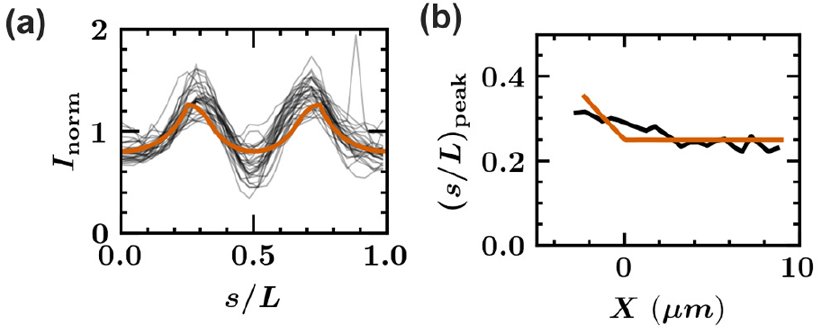
Comparison of the experimental lamin A/C spatial distribution to the minimal model predictions in the aspiration experiment. (a) Lamin A/C intensity signal measured along the curvilinear abscissa *s* at *X*=0 *µ*m. We plot the individual intensity profiles (*grey lines*) of a number of *N* = 30 cells for which we possess images at *X* = 0*±*0.3*µm*. On top is plotted the minimal model prediction (*bold orange line*). (b) Position of the fluorescence intensity peak (*s/L*)_peak_ as a function of nucleus position for different constriction sizes in the aspiration device. Orange line corresponds to the model prediction and black line corresponds to the experimentally measured average position of the first peak.

The agreement between the model predictions and experiments (Fig. 7, 8, S1 and S2) shows that, for both active and passive cell movement through confinements, the changes in lamin A/C fluorescence intensity can be well described by simple mass conservation arguments.

## 4 Discussion

We show that during both cell migration and cell aspiration, the lamin A/C fluorescence signal increases where the nucleus is compressed. This similarity suggests that in both processes, the lamin network responds in the same way even though the time scales of these two processes differ by more than two orders of magnitude and the cell behaviour is different in these two processes: in the aspiration device the cell is a passive object that is pushed through the constriction by an external pressure, while in the migration device the cell is actively involved in translocating the nucleus through the constriction. Another indication that active cell movement is not required for the observed lamin hyperfluorescence was given by the fact that the peak position of the lamin intensity profile does not depend on the migration direction, therefore, the symmetry of the hyperfluorescence indicates that this phenomenon is purely geometric. We show that the lamin hyperfluorescence is not the result of the recruitment of mobile lamins at the compression site. Instead, the hyperfluorescence at the side can be explained by simple mass conservation arguments as we illustrate with our minimal model that quantitatively reproduces the experimental lamin intensity profiles in both the migration and the aspiration device. Together, our results imply that the lamin network displays a purely passive response to both active or passive cell movement through confinements.

To keep our description as general as possible, we do not specify how the lamin network deforms exactly on length scales smaller than the (compressed) nucleus width. An interesting next step would be to elucidate what happens on these smaller scales because this could provide valuable information on the (elastic) properties of the lamin network. A possible option is that the compression by the pillars results in an increase in lamin density at the nanoscale. Another possibility is that the compression induces the wrinkling of the lamin network. In this way, the apparent surface area of the lamin network decreases without compressing the lamin on the nanoscale. In fact, this buckling has been observed for synthetic capsules flowing through constrictions: they form wrinkles on their top when the the constriction is smaller than 22 % of the non-deformed capsule diameter.[26] In addition, earlier observations on nuclei in constrictions reveal buckling of the lamin network at the side of the nucleus.[18, 19] However, we did not observe this buckling (Fig. S3). Still, we cannot exclude that wrinkling takes place. Maybe the wrinkles have a short wavelength and a small amplitude and are therefore difficult to resolve with conventional light microscopy techniques. These short wrinkling wavelengths would mean that the bending energy of the lamin network is low compared to its stretching energy.[27] Superresolution techniques might be used to assess whether short wavelength wrinkles are being formed. The change in lamin density might be tested with a recently developed lamin strain biosensor,[28] which can probe changes in lamin density at the nanoscale.

We focus on the lamin network in this paper, but the concepts that we introduce can also apply to elucidate the roles of other nuclear proteins. First, the same analysis method can be used to quantify the distribution of other proteins in the lamin shell or nuclear envelope, like nesprin-2,[19] emerin[29] and the lamina-associated-peptides.[30, 31] In addition, our quantitative description of the nuclear deformation can help to separate the contribution of this deformation from other processes that happen during confined cell migration. Since the model relies on the simple principle of mass conservation, it can easily be adapted to model other constriction geometries as we illustrate here by modelling the different geometries of the migration and aspiration device. In this way, the concepts introduced in this paper can help to further unravel the intricate mechanisms of cell migration and deformation through confinement.

## 5 Material and Methods

### 5.1 Cell Culture

Mouse Embryonic Fibroblasts (MEFs) were CRISPR-modified to create a new cell line: MEFs SYNE2-GFP LMNA-mCh and MEFs LMNA-mCh as described and validated in [19]. They express nesprin-2 giant with a green fluorescent protein (GFP) sequence and lamin A/C with a red fluorescent protein (mCherry) sequence or just lamin A/C with a red fluorescent protein (mCherry) alone. Cells are cultured at 37°C in a humidified incubator with 5% CO_2_, in DMEM (Dulbecco’s Modified Eagle Medium - Gibco) supplemented with 10% (v/v) Fetal Bovine Serum (FBS – Gibco).

### 5.2 Microfluidic devices

The migration device [18] and the aspiration device [22] have been previously described and validated. The epoxy molds (R123/R614 - Soloplast) we used were replicated from polydimethylsiloxane (PDMS) imprinted pieces coming from the lab of Jan Lammerding (Cornell University, USA). A mix of PDMS (using a 10:1 ratio polymer:crosslinker) is placed in vacuum desiccator for 20 minutes to eliminate air bubbles, then poured into one of the epoxy molds and let to cure for 4 hours in a 60°C oven. Imprinted PDMS pieces are cut using a scalpel and biopsy punches (2mm and 5mm in diameter). Glass coverslips are soaked overnight in a 0.1 M HCl solution and rinsed with H_2_O and ethanol and dried with Kimwipes. To form a microfluidic device, an imprinted PDMS piece and a treated glass coverslip are placed in a plasma cleaner for 1 minute and gently put in contact together. This process creates covalent bonds between the PDMS and the glass.[32] Devices are then directly put on a 100°C hot plate for 5 minutes to improve adhesion.

### 5.3 Cell migration experiment

Migration devices are rinsed under a microbiological safety cabinet: first once with ethanol (∼250 µL) then twice with phosphate-buffered saline (PBS - Gibco) and twice with DMEM supplemented with 10% (v/v) FBS. Cells are suspended at a concentration of 10 millions per mL in DMEM supplemented with 10% (v/v) FBS. They are seeded in the device by adding 5 µL of the suspended solution in one of the two small ports of the device. After 6 hours, enough cells are in the constricted region of the device and image acquisition can start. For that, cell medium is changed to DMEM without phenol red and with HEPES (15 mM) (Gibco), supplemented with 10% FBS (Gibco), 100 ug/mL penicillin, and 100 µg/mL streptomycin (Life Technologies).

### 5.4 Cell aspiration experiment

Immediately after assembly of the PDMS piece with the glass coverslip (see subsection 5.2), the device is filled with 25 µL of filtered 2% Bovine Serum Albumine (BSA) in PBS solution through the upper hole and put on ice until acquisition. For the acquisition, a Fluigent device is used to apply the corresponding pressures to the aspiration device. A 2 mL tube (Eppendorf), containing a suspended cells at a 5 million per mL concentration in a 2% BSA + 0.2% FBS in PBS solution, is connected to the instrument and to the upper port of the aspiration device with flexible tubes. In the same way, a 2 mL tube (Eppendorf), containing 1 mL of a 2% BSA + 0.2% FBS in PBS solution, is connected to the instrument and to the lower port of the aspiration device. In the third port of the device, the middle one, a flexible tube is placed, large enough for a pipette tip to enter. The device is first filled with cells with a pressure at the upper port tube of 100 mbar, and trapped air bubbles are removed by increasing this pressure up to 300 mbar. Finally the pressure of the upper part tube is set to 69 mbar, while the pressure in the lower part tube is set to 14 mbar. Cells are ejected from the channels at the beginning of each acquisition by using a classic lab pipette to apply a burst in pressure in the middle port.

### 5.5 FRAP experiments

For the fluorescence recovery after photobleaching (FRAP) experiments, MEFs LMNA-mCh were seeded in the migration device following the same procedure as described in section 5.3. FRAP experiments were started 16 h after seeding the cells.

### 5.6 Image acquisition

#### 5.6.1 Migration

Timelapse acquisitions are performed on an epifluorescence microscope (Nikon Ti-E) equipped with a sCMOS camera (2048 ORCA Flash 4.0 V2, Hamamatsu or Prime BSI, Teledyne), a perfect focus system, a 60x oil objective (Nikon), and a temperature and gas control chamber (set on 37°C, air at 5% CO_2_). Images are taken every 10 minutes.

#### 5.6.2 Aspiration

Timelapse acquisitions are performed on an epifluorescence microscope (Nikon Ti-E) equipped with a sCMOS camera (Prime BSI, Teledyne), a perfect focus system, a 60x oil objective (Nikon). Images are taken every second.

#### 5.6.3 FRAP

The FRAP experiments were performed on an inverted spinning disk microscope (Nikon Eclipse Ti-E/CSU-X1 Yokogawa) equipped with a 60x objective, a 561 nm laser, a FRAP module and a thermostatic chamber. Nuclei that were inside a constriction were selected for FRAP experiments. The bleach region was a 16 *µ*m by 4 *µ*m rectangle spanning the width of the constriction. The bleaching was obtained by scanning this region 20 times at a 100 % laser intensity. 60 images were recorded before bleaching and 600 images were recorded after bleaching. The image acquisition time interval was 1 s.

### 5.7 Image analysis

Movies are analysed using Image J/Fiji and Python. The projected nucleus surface is detected with Image J/Fiji by using the ”analyse particles” function on a threshold (median filter to 5.0 radius, normalised by 0.4% and autolocal threshold ”Bernsen” 5) applied on the mCherry image (corresponding to the lamin A/C signal). The nucleus contour is taken as a 1 µm wide band at the edge of the nucleus. The contour is divided into 30 segments along the curvilinear abscissa *s* and the fluorescence intensity inside each segment is measured. *s* = 0 corresponds to the back of the nucleus. It is defined as the back coordinate of the longest axis of the nucleus that passes through the barycenter of the nucleus. The contour coordinate *s* is normalised by dividing by the total contour length *L*. The fluorescence intensity in the contour is corrected for the background intensity by subtracting the average fluorescence intensity in 5 µm by 5 µm square region inside the PDMS pillar. Subsequently, this corrected contour fluorescence intensity is normalised by dividing by the average intensity in the contour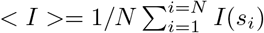.

#### 5.7.1 Origin points and nucleus position

In the migration device, the origin of the *x* axis is set at the center of the constriction pillar. In the aspiration device, the origin of the *x* axis is set at the entry of the channel (Fig. 1).

In both migration and aspiration, the position of a nucleus is defined by its surface geometrical barycenter (*X, Y* ), specifically 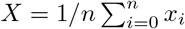 and *Y* = 1*/n* Σ _*i*_*y*_*i*_ with (*x*_*i*_, *y*_*i*_) the coordinates of each pixel *i* of the nucleus surface and *n* the number of pixels in the nucleus surface.

#### 5.7.2 Peak position determination

An intensity profile is used for the peak position determination if the standard deviation of its intensity values is larger than 0.05. In that case, the intensity profile is separated into two parts: a first one for 0 ≤ *s/L <* 0.5 and a second one for 0.5 ≤ *s/L* ≤ 1. For both parts, the value *s/L* at which the intensity has its maximum is taken as peak position. In this way, we obtain the peak positions for many intensity profiles each with their own nucleus position *X*. Subsequently, we bin the calculated peak positions according to their nucleus position to calculate the average peak position as function of the nucleus position *X*. The used bin size is Δ*X* = 2 *µ*m for the migration experiments and Δ*X* = 0.5 *µ*m for the aspiration experiments.

## Author Contributions

CS conceived the project. SA and IB performed experimental work and analysis. ER performed the modelling. All authors co-wrote the manuscript.

## Acknowledgements

We thank the members of the Breast Cancer Biology Group, Institut Curie - PSL Research University, Paris, France for their help and fruitful discussions. We thank Julie Plastino and Basile Audoly for many fruitful discussions, and Hasti Honari for accompanying SA in setting up the micropipette experiment at LPENS. This work was supported by an institutional Institut Curie grant and by the French Agence Nationale de la Recherche (ANR), under grant ANR-22-CE30-0030 (project SQUEEZACTIV). It was partly carried out at the Cell and Tissue Imaging (PICT-IBiSA) facility, Institut Curie, member of the French National Research Infrastructure France-BioImaging (ANR10-INBS-04). The work of IB was supported by the Dutch Research Council (NWO) through the Rubicon programme (project number 019.221EN.001).

## References

(1) Aman, A.; Piotrowski, T. Cell migration during morphogenesis. Dev. Biol. 2010, 341, 20–33.

(2) Smith, L. R.; Irianto, J.; Xia, Y.; Pfeifer, C. R.; Discher, D. E. Constricted migration modulates stem cell differentiation. Mol. Biol. Cell 2019, 30, 1985–1999.

(3) Weninger, W.; Biro, M.; Jain, R. Leukocyte migration in the interstitial space of non-lymphoid organs. Nat. Rev. Immunol 2014, 14, 232–246.

(4) Denais, C. M.; Gilbert, R. M.; Isermann, P.; McGregor, A. L.; Te Lindert, M.; Weigelin, B.; Davidson, P. M.; Friedl, P.; Wolf, K.; Lammerding, J. Nuclear envelope rupture and repair during cancer cell migration. Science 2016, 352, 353–358.

(5) Wolf, K.; Te Lindert, M.; Krause, M.; Alexander, S.; Te Riet, J.; Willis, A. L.; Hoffman, R. M.; Figdor, C. G.; Weiss, S. J.; Friedl, P. Physical limits of cell migration: control by ECM space and nuclear deformation and tuning by proteolysis and traction force. J. Cell Biol 2013, 201, 1069–1084.

(6) McGregor, A. L.; Hsia, C.-R.; Lammerding, J. Squish and squeeze—the nucleus as a physical barrier during migration in confined environments. Curr. Opin. Cell Biol. 2016, 40, 32–40.

(7) Kalukula, Y.; Stephens, A. D.; Lammerding, J.; Gabriele, S. Mechanics and functional consequences of nuclear deformations. Nat. Rev. Mol. Cell Biol. 2022, 23, 583–602.

(8) Niethammer, P. Components and mechanisms of nuclear mechanotransduction. Annu. Rev. Cell Dev. Biol. 2021, 37, 233–256.

(9) Turgay, Y.; Eibauer, M.; Goldman, A. E.; Shimi, T.; Khayat, M.; Ben-Harush, K.; Dubrovsky-Gaupp, A.; Sapra, K. T.; Goldman, R. D.; Medalia, O. The molecular architecture of lamins in somatic cells. Nature 2017, 543, 261–264.

(10) Kittisopikul, M.; Virtanen, L.; Taimen, P.; Goldman, R. D. Quantitative analysis of nuclear lamins imaged by super-resolution light microscopy. Cells 2019, 8, 361.

(11) Lammerding, J.; Fong, L. G.; Ji, J. Y.; Reue, K.; Stewart, C. L.; Young, S. G.; Lee, R. T. Lamins A and C but not lamin B1 regulate nuclear mechanics. J. Biol. Chem. 2006, 281, 25768–25780.

(12) Zwerger, M.; Roschitzki-Voser, H.; Zbinden, R.; Denais, C.; Herrmann, H.; Lammerding, J.; Grütter, M. G.; Medalia, O. Altering lamina assembly reveals lamina-dependent and-independent functions for A-type lamins. J. Cell Sci. 2015, 128, 3607–3620.

(13) Worman, H. J.; Bonne, G. “Laminopathies”: A wide spectrum of human diseases. Exp. Cell Res. 2007, 313, 2121–2133.

(14) Leong, E. L.; Khaing, N. T.; Cadot, B.; Hong, W. L.; Kozlov, S.; Werner, H.; Wong, E. S. M.; Stewart, C. L.; Burke, B.; Lee, Y. L. Nesprin-1 LINC complexes recruit microtubule cytoskeleton proteins and drive pathology in Lmna-mutant striated muscle. Hum. Mol. Genet. 2023, 32, 177– 191.

(15) Chiarini, F.; Paganelli, F.; Balestra, T.; Capanni, C.; Fazio, A.; Manara, M. C.; Landuzzi, L.; Petrini, S.; Evangelisti, C.; Lollini, P.-L., et al. Lamin A and the LINC complex act as potential tumor suppressors in Ewing Sarcoma. Cell Death & Disease 2022, 13, 346.

(16) Dubik, N.; Mai, S. Lamin A/C: Function in normal and tumor cells. Cancers 2020, 12, 1–21.

(17) Bell, E. S.; Shah, P.; Zuela-Sopilniak, N.; Kim, D.; Varlet, A.-A.; Morival, J. L.; McGregor, A. L.; Isermann, P.; Davidson, P. M.; Elacqua, J. J., et al. Low lamin A levels enhance confined cell migration and metastatic capacity in breast cancer. Oncogene 2022, 41, 4211–4230.

(18) Davidson, P. M.; Sliz, J.; Isermann, P.; Denais, C.; Lammerding, J. Design of a microfluidic device to quantify dynamic intra-nuclear deformation during cell migration through confining environments. Integr. Biol. 2015, 7, 1534–1546.

(19) Davidson, P. M.; Battistella, A.; Déjardin, T.; Betz, T.; Plastino, J.; Borghi, N.; Cadot, B.; Sykes, C. Nesprin-2 accumulates at the front of the nucleus during confined cell migration. EMBO Rep. 2020, 21, 1–14.

(20) Kim, D.-H.; Li, B.; Si, F.; Phillip, J. M.; Wirtz, D.; Sun, S. X. Volume regulation and shape bifurcation in the cell nucleus. J. Cell Sci. 2015, 128, 3375–3385.

(21) Cao, X.; Moeendarbary, E.; Isermann, P.; Davidson, P. M.; Wang, X.; Chen, M. B.; Burkart, A. K.; Lammerding, J.; Kamm, R. D.; Shenoy, V. B. A Chemomechanical Model for Nuclear Morphology and Stresses during Cell Transendothelial Migration. Biophys. J. 2016, 111, 1541–1552.

(22) Davidson, P. M.; Fedorchak, G. R.; Mondésert-Deveraux, S.; Bell, E. S.; Isermann, P.; Aubry, D.; Allena, R.; Lammerding, J. High-throughput microfluidic micropipette aspiration device to probe time-scale dependent nuclear mechanics in intact cells. Lab Chip 2019, 19, 3652–3663.

(23) Broers, J. L.; Machiels, B. M.; Van Eys, G. J.; Kuijpers, H. J.; Manders, E. M.; Van Driel, R.; Ramaekers, F. C. Dynamics of the nuclear lamina as monitored by GFP-tagged A-type lamins. J. Cell Sci. 1999, 112, 3463–3475.

(24) Moir, R. D.; Yoon, M.; Khuon, S.; Goldman, R. D. Nuclear lamins A and B1: Different pathways of assembly during nuclear envelope formation in living cells. J. Cell Biol. 2000, 151, 1155–1168.

(25) Naetar, N.; Ferraioli, S.; Foisner, R. Lamins in the nuclear interior-life outside the lamina. J. Cell Sci. 2017, 130, 2087–2096.

(26) Dawson, G.; Häner, E.; Juel, A. Extreme deformation of capsules and bubbles flowing through a localised constriction. Procedia IUTAM 2015, 16, 22–32.

(27) Cerda, E.; Mahadevan, L. Geometry and Physics of Wrinkling. Phys. Rev. Lett. 2003, 90, 074302.

(28) Danielsson, B. E.; George Abraham, B.; Mäntylä, E.; Cabe, J. I.; Mayer, C. R.; Rekonen, A.; Ek, F.; Conway, D. E.; Ihalainen, T. O. Nuclear lamina strain states revealed by intermolecular force biosensor. Nat. Commun. 2023, 14, 3867.

(29) Lavenus, S. B.; Vosatka, K. W.; Caruso, A. P.; Ullo, M. F.; Khan, A.; Logue, J. S. Emerin regulation of nuclear stiffness is required for fast amoeboid migration in confined environments. J. Cell Sci. 2022, 135, jcs259493.

(30) Chen, N. Y.; Kim, P. H.; Tu, Y.; Yang, Y.; Heizer, P. J.; Young, S. G.; Fong, L. G. Increased expression of LAP2β eliminates nuclear membrane ruptures in nuclear lamin–deficient neurons and fibroblasts. Proc. Natl. Acad. Sci. U.S.A 2021, 118, e2107770118.

(31) Jung-Garcia, Y.; Maiques, O.; Monger, J.; Rodriguez-Hernandez, I.; Fanshawe, B.; Domart, M.-C.; Renshaw, M. J.; Marti, R. M.; Matias-Guiu, X.; Collinson, L. M., et al. LAP1 supports nuclear adaptability during constrained melanoma cell migration and invasion. Nat. Cell Biol. 2023, 25, 108–119.

(32) Borók, A.; Laboda, K.; Bonyár, A. PDMS bonding technologies for microfluidic applications: A review. Biosensors 2021, 11, 292.

