## Supporting Information for "The nuclear lamin network passively responds to both active or passive cell movement through confinements"

### S1 Model and experiment comparisons

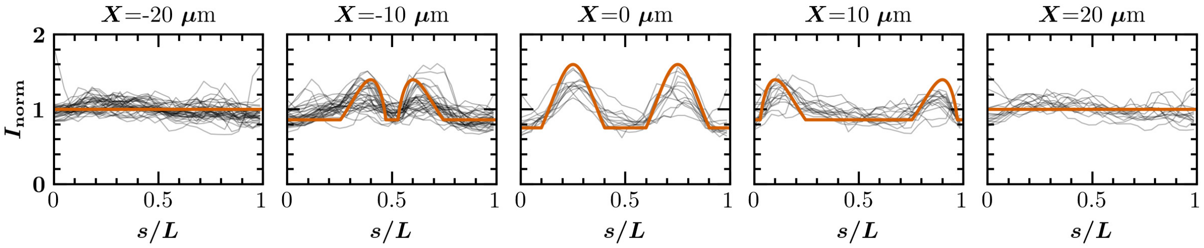


**Figure S1.** Comparison of the lamin A/C distribution with the minimal model predictions for different positions in a 3 $\mu$m constriction in a migration device ($X$= -20 $\mu$m, $X$ = -10 $\mu$m, $X$ = 0 $\mu$m, $X$ = 10 $\mu$m and $X$ = 20 $\mu$m).

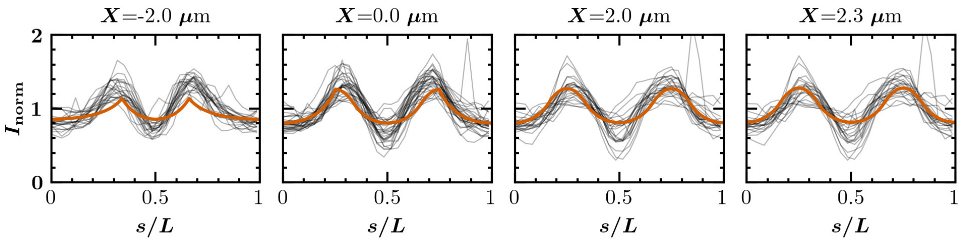


**Figure S2.** Comparison of the lamin A/C distribution with the minimal model predictions for different positions in the aspiration device ($X$= -3 $\mu$m, $X$ = 0 $\mu$m, $X$ = 3 $\mu$m and $X$ = 5.5 $\mu$m).

‡These authors contributed equally to this work

### S2 Z-stack microscopy measurements

Z-stack acquisitions of MEFS Lmna mCh in a migration device were performed on an inverted spinning disk microscope (Nikon Eclipse Ti-E/CSU-X1 Yokogawa) equipped with a sCMOS camera (Prime BSI, Teledyne), a 60x CFI Plan Apo IR SR water immersion objective (Nikon), a 561 nm laser and a temperature and gas control chamber (set on 37 °C, air at 5% $CO_2$). For the z-stack measurements, 26 stack images were taken with a height interval of $\Delta z = 0.4\mu$m.

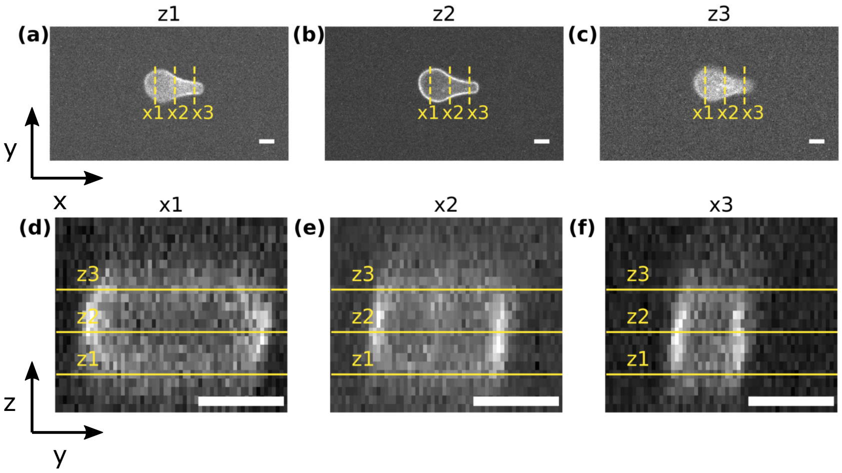


**Figure S3.** Example spinning disk lamin A/C fluorescence images of a nucleus in a 3 $\mu$m constriction. Example images at the bottom (z1 (a)), middle (z2,(b)) and top (z3,(c)) of the cell are shown. Vertical dashed line indicate the positions where vertical cross sections are taken, shown in (d) for position x1, in (e) for position x2 and in (f) for position x3. The horizontal yellow lines in these cross sections (d-f) indicate the heights z1, z2 and z3 at which the example images of (a-c) are taken. In all image, the scale bar = 4 $\mu$m.
